# Increased ketohexokinase-A governs fructose-induced podocyte hypertrophy by IL-6/STAT3 signaling activation

**DOI:** 10.1101/2020.12.28.424520

**Authors:** Jie Zhou, Dong-Mei Zhang, Jie Yang, Hong Ding, Tu-Shuai Li, Zhi-Hong Liu, Li Chen, Rui-Qing Jiao, Ling-Dong Kong

## Abstract

Glomerular hypertrophy is crucial for podocyte damage and proteinuria. Our previous study showed that fructose induced podocyte injury. However, the molecular mechanism underlying podocyte hypertrophy under fructose is unclear. We observed that fructose significantly initiated the hypertrophy in rat glomeruli and cultured differentiated human podocytes (HPCs). Consistently, it induced inflammatory response with the down-regulation of zinc-finger protein tristetraprolin (TTP) and the activation of interleukin-6 (IL-6)/signal transducer and activator of transcription 3 (STAT3) signaling in these animal and cell models. Subsequently, high-expression of miR-92a-3p and its target protein cyclin-dependent kinase inhibitor p57 (P57) down-regulation, representing the abnormal proliferation and apoptosis, were observed *in vivo* and *in vitro*. Moreover, fructose increased ketohexokinase-A (KHK-A) in rat glomeruli and HPCs. Animal-free recombinant human IL-6, maslinic acid and *TTP* siRNA were used to manifest that fructose may decrease TTP to activate IL-6/STAT3 signaling in podocyte overproliferation and apoptosis, causing podocyte hypertrophy. *KHK-A* siRNA transfection further demonstrated that the inactivation of IL-6/STAT3 to relieve podocyte hypertrophy mediated by inhibiting KHK-A to increase TTP may be a novel strategy for fructose-associated podocyte injury and proteinuria.

**Graphic Abstract:** 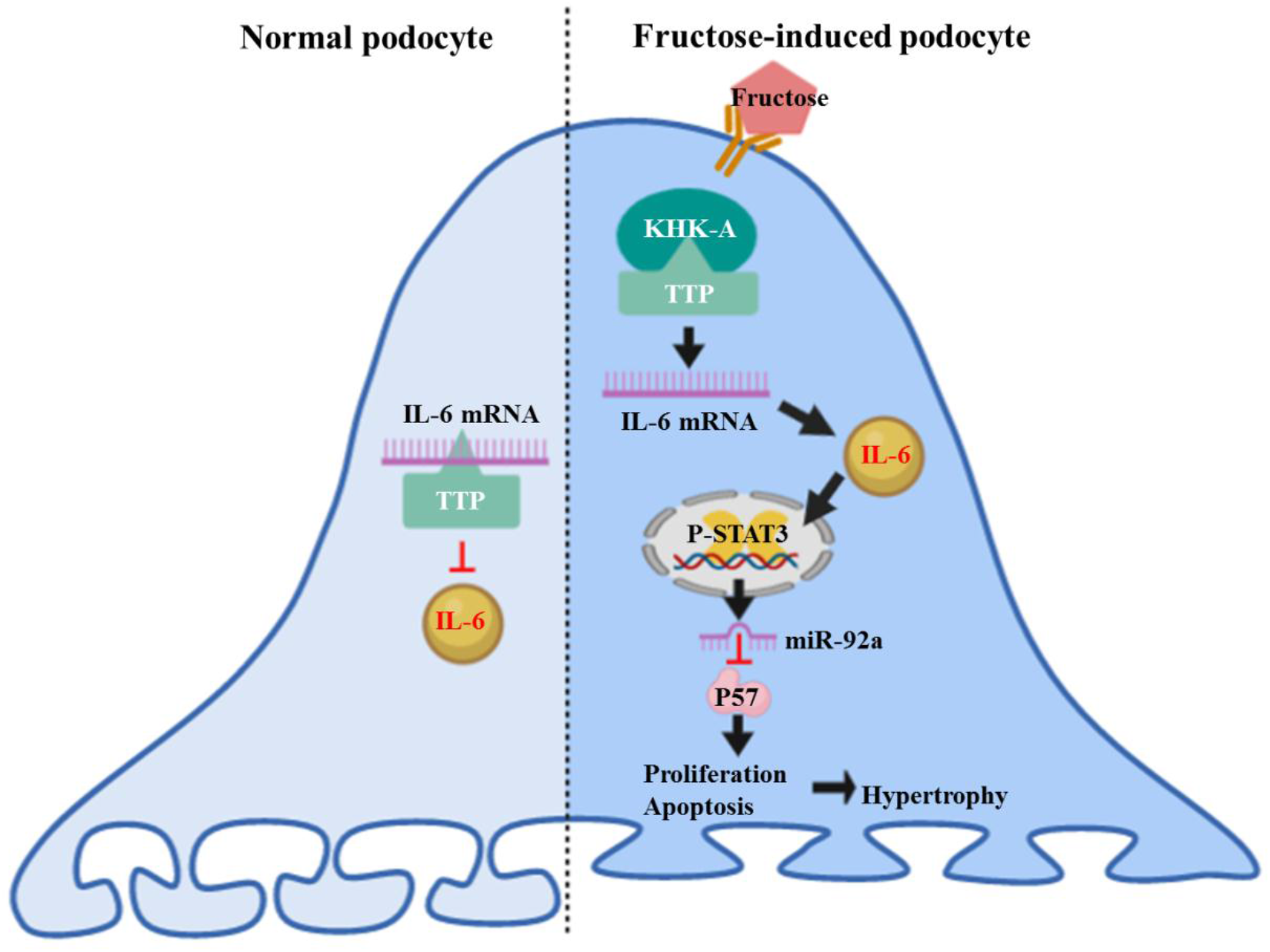

## Introduction

The glomerular filtration barrier is constructed by podocytes, fenestrated endothelial cells and glomerular basement membrane (GBM) (Tian et al., 2014). Glomerular hypertrophy makes podocytes sensitive to increased glomerular filtration pressure, inducing podocyte instability in patients with type 2 diabetes and focal segmental glomerulosclerosis (Lemley et al., 2000; Puelles et al., 2019). Glomerular podocytes, terminal differentiated cells with limited self-renewal ability, are difficult to repair and regenerate after damage (Mulay et al., 2013). High fructose diet is reported to increase kidney size in rats (Sánchez-Lozada et al., 2007). Our previous study showed that high fructose feeding induced glomerular injury with podocyte apoptosis and inflammation in rats (Wang et al., 2015; Li et al., 2019). However, whether high fructose causes glomerular and podocyte hypertrophy remains unclear.

Interleukin-6 (IL-6) is a pleiotropic cytokine (Feigerlová and Battaglia-Hsu, 2017). IL-6 overexpression and podocyte hypertrophy are observed in adriamycin-induced nephropathy mice (Mulay et al., 2013). In cultured mouse podocytes, high glucose induces the secretion of IL-6 with cell hypertrophy and apoptosis (Jo et al., 2016; Shi et al., 2016). Addition of exogenous recombinant IL-6 is reported to activate signal transducer and activator of transcription 3 (STAT3) in primary cultured podocytes (Henique et al., 2017). Activation of IL-6/STAT3 signaling is detected in angiotensin II-induced hypertrophic cardiomyopathy mice with cardiac inflammation and hypertrophy (Chen et al., 2019). These observations indicate that IL-6/STAT3 activation may promote podocyte hypertrophy.

Tristetraprolin (TTP), known as an anti-inflammatory factor, can mediate the degradation of mRNAs of inflammatory factors (Guo et al., 2020). TTP expression is obviously decreased, accompanied with elevated IL-6 level in mouse kidney of doxorubicin-induced nephrotic syndrome (Zhang et al., 2020). Its down-regulation is negatively correlated with IL-6 expression in diabetic patients with proteinuria (Liu et al., 2015).

It is known that ketohexokinase (KHK) promotes fructose catabolism (Hayasaki et al., 2019). Silence of KHK expression by lentiviral transfection causes significant reduction of IL-6 production in glucose-exposed human proximal tubular epithelial cells (Lanaspa et al., 2014). KHK-A is primarily expressed in kidney (Diggle et al., 2009). Thus, we speculated that KHK-A dysregulation in respond to fructose exposure may suppress TTP expression to activate IL-6/STAT3 signaling in podocytes. Of note, microRNA-92a (miR-92a) is one of the target genes of STAT3 (Henique et al., 2017). Its up-regulation is detected in the podocytes of rapidly progressive glomerulonephritis patients with IL-6/STAT3 activation (Henique et al., 2017). Cyclin-dependent kinase inhibitor p57 (P57) leads to G1 phase cell cycle arrest (Yan et al., 1997). MiR-92a is overexpressed in the glomeruli of nephrotoxic nephritis mice, and its down-regulation increases P57 expression and limits podocyte proliferation in primary podocytes transfected with anti-miR-92a (Henique et al., 2017). Overexpression of miR-92a-3p increases cell overproliferation in human renal cancer cells (Zeng et al., 2020). Therefore, the investigation of mechanism of podocyte overproliferation and apoptosis may provide a deeper insight to IL-6/STAT3 signaling activation in fructose-induced podocyte hypertrophy.

In this study, we demonstrated that high fructose induced hypertrophy in rat glomeruli and differentiated human podocytes (HPCs). Meanwhile, the activation of IL-6/STAT3 signaling and down-regulation of TTP was observed in these animal and cell models with podocyte overproliferation and apoptosis. Interfering with KHK-A could promote TTP expression to inhibit the activation of IL-6/STAT3 signaling in fructose-exposed HPCs. TTP low-expression to activate IL-6/STAT3 was sufficient to up-regulate miR-92a-3p, down-regulate P57 to induce podocyte overproliferation and apoptosis in fructose-induced podocyte hypertrophy. These results suggested that IL-6/STAT3 signaling activation exacerbates fructose-induced podocyte hypertrophy by increasing KHK-A to down-regulate TTP.

## Results

### High fructose induces glomerular hypertrophy in rats and podocyte hypertrophy in differentiated HPCs, with podocyte overproliferation and apoptosis

As expected (Wang et al., 2015), glomerular podocyte injury was observed in fructose-fed rats with the increases of body weight, serum levels of uric acid and creatinine, as well as urea nitrogen level and proteinuria at week 16th (Table 2, Supplementary Fig. 1). At week 12th and 16th, the mean size of glomeruli from fructose-fed rats was significantly larger than that from normal animals (Fig. 1A). Fructose-induced podocyte hypertrophy was also observed in differentiated HPCs (Fig. 1B). EdU-positive podocytes were increased remarkably in fructose-exposed differentiated HPCs (Fig. 1C), while KI67-stained podocytes were also augmented in the glomeruli of fructose-fed rats (Fig. 1D), confirming podocyte overproliferation. Moreover, fructose increased the proportion of podocytes in S phase (Fig. 1E). Consistently, fructose-induced podocyte apoptosis was detected in differentiated HPCs by Flow cytometry analysis (Fig. 1F) and rat glomeruli by TUNEL analysis (Fig. 1G). These results demonstrated that high fructose induced glomerular podocyte hypertrophy, as well as podocyte overproliferation and apoptosis in these animal and cell models.

**Fig.1.**
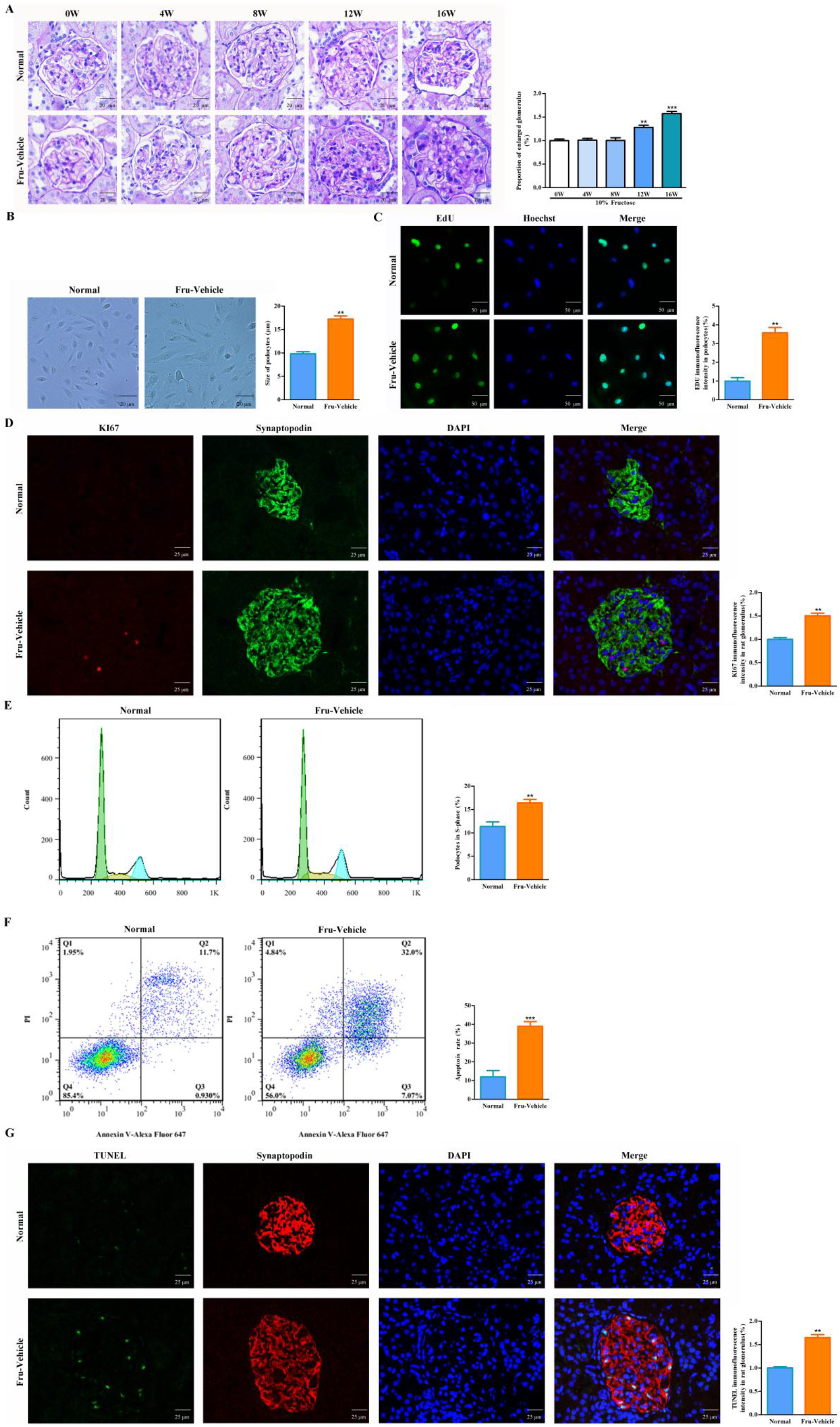
Fructose induces glomerular hypertrophy in rats and podocyte hypertrophy in differentiated HPCs, with podocyte overproliferation and apoptosis *in vivo* and *in vitro*. (**A**) Periodic acid Schiff (PAS) staining showing development of glomerular lesions after fructose intake for 0, 4, 8, 12, 16 weeks (scale bar, 20 μm, n = 3). (**B**) Podocyte size in differentiated HPCs cultured with or without 5 mM fructose for 72 h was measured respectively (scale bar, 20 μm, n = 3). (**C**) Representative immunofluorescence showing increased EDU staining in differentiated HPCs cultured with or without 5 mM fructose for 72 h (scale bar, 50 μm, n = 3). (**D**) Representative immunofluorescence showing increased KI67 staining in podocytes of fructose-fed rat kidney glomeruli (scale bar, 25 μm, n = 3). (**E**) Flow cytometry analysis of cell cycle in differentiated HPCs cultured with or without 5 mM fructose for 72 h (n = 6), respectively. (**F**) Flow cytometry analysis of apoptosis in differentiated HPCs cultured with or without 5 mM fructose for 72 h (n = 6), respectively. (**G**) Representative TUNEL staining showing apoptotic cells occurred in the kidney glomeruli of fructose-fed rats (scale bar, 25 μm, n = 3). All data are expressed as mean ± SEM. ** *p* < 0.01, *** *p* < 0.001 compared with normal animal control group or normal cell control.

### High fructose suppresses TTP to activate IL-6/STAT3 signaling in podocyte hypertrophy

In this study, a significant decrease of TTP protein levels was detected in the glomeruli of fructose-fed rats and fructose-exposed differentiated HPCs (Fig. 2A-B). High fructose increased IL-6 levels in serum and renal cortex of rats (Fig. 2C), as well as in the culture media and lysate of differentiated HPCs (Fig. 2D). IL-6 mRNA levels were increased in fructose-fed rat glomeruli and fructose-exposed differentiated HPCs (Fig. 2E-F). InBio-Discover analysis revealed that IL-6 overproduction may probably correlate with STAT3 signaling activation (Fig. 2G). Besides, fructose significantly increased STAT3 phosphorylation in rat glomeruli (Fig. 2H) and differentiated HPCs (Fig. 2I). Furthermore, exogenous IL-6 incubation increased STAT3 phosphorylation in differentiated podocytes (Fig. 2J-K). Decreased protein levels of nephrin and podocin, as well as podocyte hypertrophy were also detected in IL-6-treated differentiated HPCs (Fig. 2L-M). Maslinic acid, a iterpenoid compound with anti-inflammatory effect (Reyes-Zurita et al., 2009), ameliorated fructose-induced IL-6 overproduction and STAT3 over-phosphorylation in differentiated HPCs (Fig. 2N-P). Therefore, the activation IL-6/STAT3 signaling induced by fructose contributed to podocyte hypertrophy.

**Fig.2.**
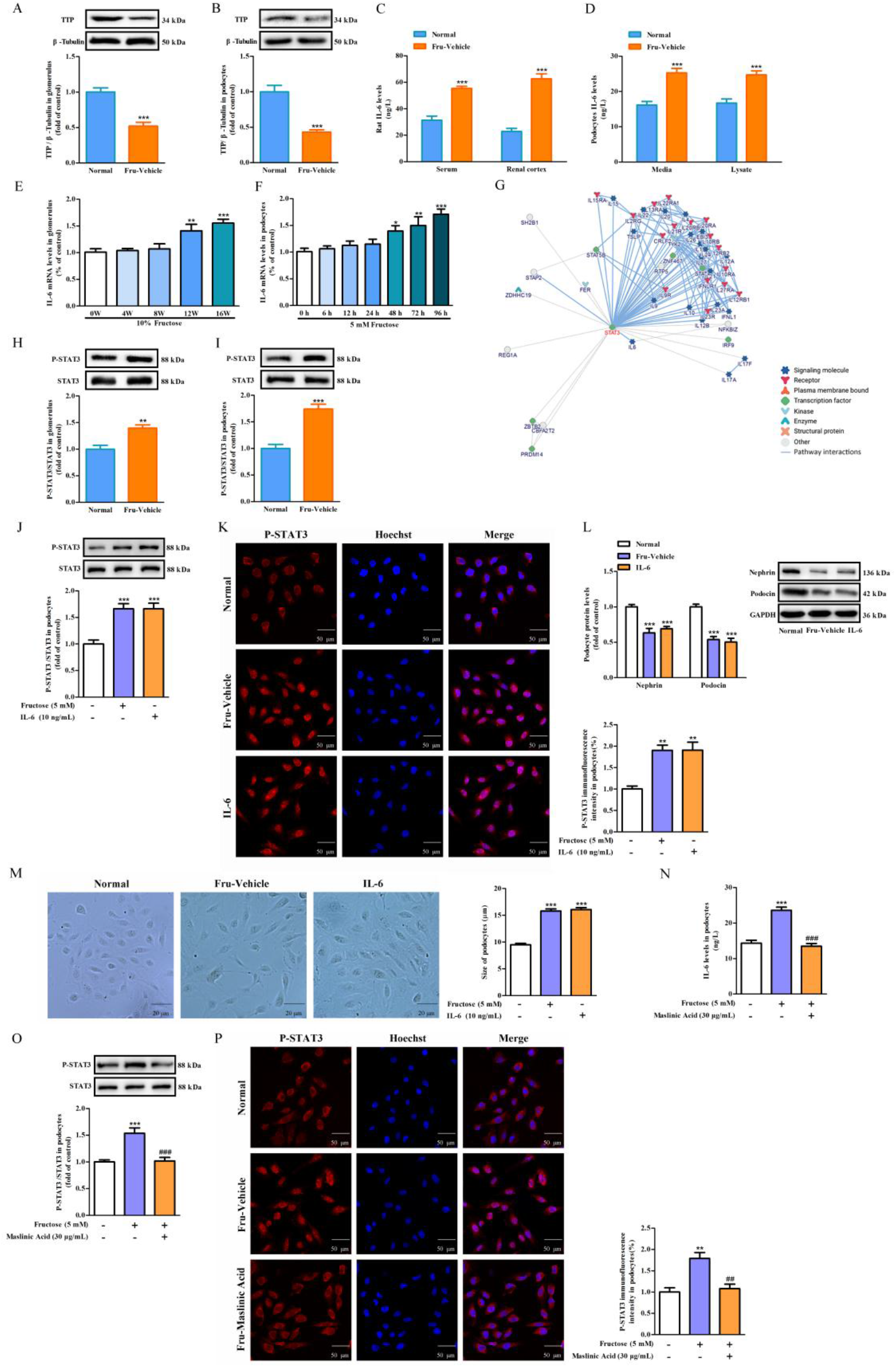
Fructose down-regulates TTP protein expression to activate IL-6/STAT3 signaling in rat kidney glomeruli and fructose-exposed differentiated HPCs. (**A-B**) Western blot analysis of TTP protein levels in the kidney glomeruli (16W) of normal control, fructose vehicle rats or differentiated HPCs cultured with or without 5 mM fructose for 72 h. Relative protein level of TTP was normalized to β-Tubulin (n = 6), respectively. (**C**) ELISA analysis of IL-6 levels (n = 6) in serum and renal cortex of rats fed with or without 10% fructose for 16W, respectively. (**D**) ELISA analysis of IL-6 levels (n = 6) in culture media and lysate in differentiated HPCs cultured with or without 5 mM fructose for 72 h, respectively. (**E-F**) qRT-PCR analysis of IL-6 levels in the kidney glomeruli (n = 6) of normal control, fructose vehicle rats or differentiated HPCs (n = 6) cultured with or without 5 mM fructose, respectively. (**G**) The protein-protein interaction of IL-6 and STAT3 was analyzed and visualized by InBio-Discover. (**H-I**) Western blot analysis of P-STAT3 protein levels in the kidney glomeruli (16W) of normal control, fructose vehicle rats or differentiated HPCs cultured with or without 5 mM fructose for 72 h. Relative protein level of P-STAT3 was normalized to STAT3 (n = 6), respectively. (**J**) Western blot analysis of P-STAT3 protein levels in differentiated HPCs incubated with or without 5 mM fructose in the presence or absence of 10 ng/mL recombinant human IL-6 for 72 h. Relative protein level of P-STAT3 was normalized to STAT3 (n = 6), respectively. (**K**) Immunofluorescence analysis of P-STAT3 (red) in differentiated HPCs cultured with or without 5 mM fructose in the presence or absence of 10 ng/mL recombinant human IL-6 for 72 h (scale bar, 50 μm, n = 3), respectively. (**L**) Western blot analysis of nephrin and podocin protein levels in differentiated HPCs incubated with or without 5 mM fructose in the presence or absence of 10 ng/mL recombinant human IL-6 for 72 h. Relative protein levels of nephrin and podocin were normalized to GAPDH (n = 6), respectively. (**M**) Podocyte size in differentiated HPCs cultured with or without 5 mM fructose in the presence or absence of 10 ng/mL recombinant human IL-6 for 72 h was measured separately (scale bar, 20 μm, n = 3). (**N**) ELISA analysis of IL-6 levels (n = 6) from lysate in differentiated HPCs cultured with or without 5 mM fructose in the presence or absence of 30 μg/mL maslinic acid for 72 h, respectively. (**O**) Western blot analysis of P-STAT3 protein levels in differentiated HPCs incubated with or without 5 mM fructose in the presence or absence of 30 μg/mL maslinic acid for 72 h. Relative protein level of P-STAT3 was normalized to STAT3 (n = 6), respectively. (**P**) Immunofluorescence analysis of P-STAT3 (red) in differentiated HPCs cultured with or without 5 mM fructose in the presence or absence of 30 μg/mL maslinic acid for 72 h (scale bar, 50 μm, n = 3), respectively. All data are expressed as mean ± SEM. * *p* < 0.05, ** *p* < 0.01, *** *p* < 0.001 compared with normal animal control group or normal cell control; ^##^*p* < 0.01, ^###^*p* < 0.001 compared with compared with fructose-vehicle animal group or fructose vehicle cell group.

Further experiments were done to investigate the correlation between TTP low expression and IL-6/STAT3 signaling activation in fructose-induced podocyte hypertrophy. Differentiated HPCs were transfected with *TTP* siRNA. TTP mRNA and protein levels were analyzed by qRT-PCR and Western blot to confirm the transfection efficiency (Fig. 3A-B). *TTP* siRNA transfection did not further induce IL-6 up-regulation and STAT3 phosphorylation in fructose-exposed differentiated HPCs (Fig. 3C-D). Consistently, podocyte hypertrophy induced by fructose was not aggravated by *TTP* siRNA transfection (Fig. 3E). These results demonstrated that high fructose may suppress TTP expression to induce podocyte hypertrophy by activating IL-6/STAT3 signaling.

**Fig.3.**
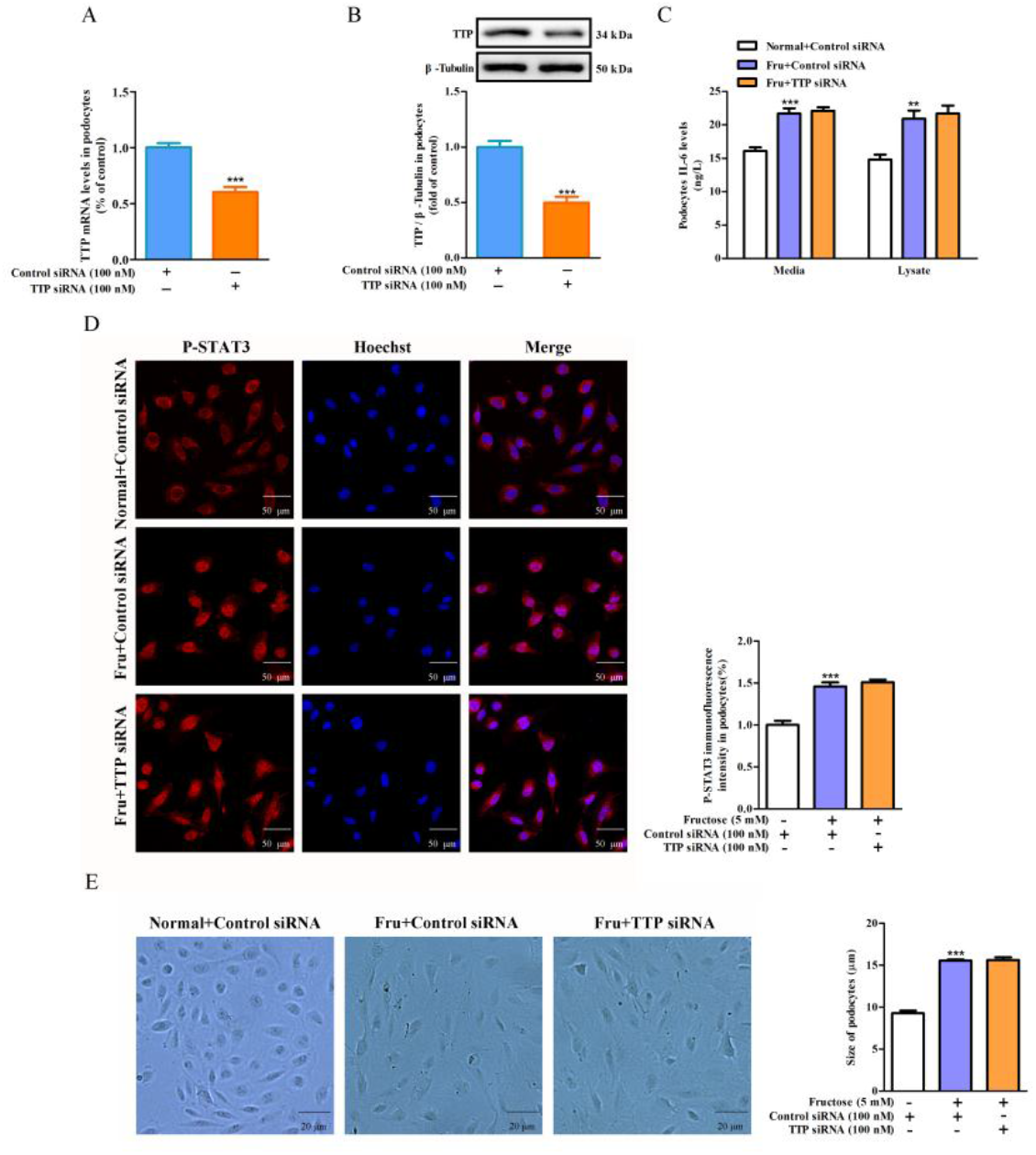
Down-regulation of TTP expression does not further activate IL-6/STAT3 signaling in fructose-exposed differentiated HPCs. (**A-B**) The transfection efficiency of *TTP* siRNA (100 nM) in differentiated HPCs and control siRNA was evaluated by qRT-PCR analysis (24 h, n = 6), and Western blot analysis (48 h, n = 6), respectively. (**C**) ELISA analysis of IL-6 levels (n = 6) from media and lysate in differentiated HPCs transfected with *TTP* siRNA or control siRNA, and then incubated with or without 5 mM fructose for 72 h, respectively. (**D**) Immunofluorescence analysis of P-STAT3 (red) in differentiated HPCs transfected with *TTP* siRNA or control siRNA, and then incubated with or without 5 mM fructose for 72 h (scale bar, 50 μm, n = 3), respectively. (**E**) Podocyte size in differentiated HPCs transfected with *TTP* siRNA or control siRNA, and then incubated with or without 5 mM fructose for 72 h was measured separately (scale bar, 20 μm, n = 3). All data are expressed as mean ± SEM. ** *p* < 0.01, *** *p* < 0.001 compared with normal animal control group or normal cell control.

### High fructose induces inflammatory response to up-regulate miR-92a-3p and reduces its target protein P57 expression in podocyte overproliferation and apoptosis

Meanwhile, increased miR-92a-3p mRNA (Fig. 4A-B) and decreased P57 (Fig. 4C-D) protein levels were detected in the glomeruli of fructose-fed rats and fructose-exposed differentiated HPCs. MiR-92a is reported to target the end of the coding sequence of P57, its down-regulation is associated with a high rate of podocyte proliferation (Henique et al., 2017). TargetScan analysis showed that the potential binding sites of has- and rno-miR-92a-3P located between 153 and 159 bp, 158 and 164 bp region at the 3’-end of P57 coding sequence, respectively (Fig. 4E). IL-6 stimulation also increased miR-92a-3p and decreased P57 protein levels in differentiated HPCs (Fig. 4F-G). Podocyte overproliferation manifested by EdU immunofluorescence staining and S-phase cell quantification (Fig. 4H-I) as well as podocyte apoptosis (Fig. 4J) were observed in IL-6-incubated HPCs. *TTP* siRNA did not further increase EdU-positive podocytes, the proportion of podocytes in S phase and podocyte apoptosis in fructose-exposed differentiated HPCs (Fig. 4K-M). These data suggested that fructose may enhance the IL-6/STAT3 signaling to induce miR-92a-3p up-regulation and P57 reduction in podocyte overproliferation and apoptosis.

**Fig.4.**
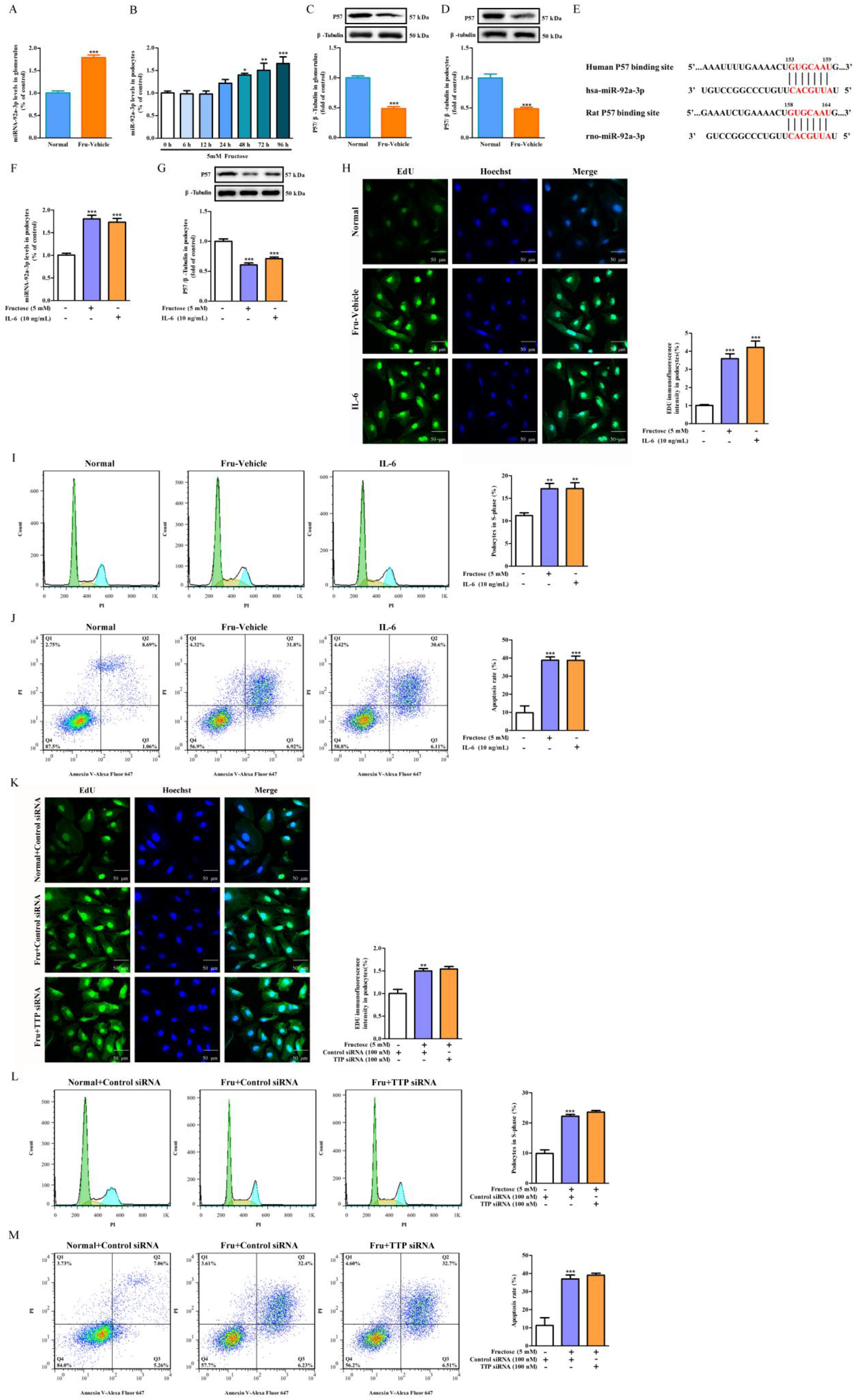
High fructose induces the activation of IL-6/STAT3 signaling to upregulate miR-92a-3p and reduced P57 expression in podocyte proliferation and apoptosis. (**A**) qRT-PCR analysis of miR-92a-3p levels (16W) in rat kidney glomeruli (n = 6) of normal control, fructose vehicle, respectively. (**B**) qRT-PCR analysis of miR-92a-3p levels (0, 6, 12, 24, 48, 72, 96 h) in differentiated HPCs (n = 6) cultured with or without 5 mM fructose, respectively. (**C-D**) Western blot analysis of P57 protein levels in the kidney glomeruli (16W) of normal control, fructose vehicle rats or differentiated HPCs cultured with or without 5 mM fructose for 72 h. Relative protein level of P57 was normalized to β-Tubulin (n = 6), respectively. (**E**) Sequence alignment between miR-92a-3p and the binding site of P57 sequence in rat and human. (**F**) qRT-PCR analysis of miR-92a-3p levels (n = 6) in differentiated HPCs incubated with or without 5 mM fructose in the presence or absence of 10 ng/mL recombinant human IL-6 for 72 h, respectively. (**G**) Western blot analysis of P57 protein levels in differentiated HPCs incubated with or without 5 mM fructose in the presence or absence of 10 ng/mL recombinant human IL-6 for 72 h. Relative protein level of P57 was normalized to β-Tubulin (n = 6), respectively. (**H**) Immunofluorescence analysis of EDU (green) in differentiated HPCs cultured with or without 5 mM fructose in the presence or absence of 10 ng/mL recombinant human IL-6 for 72 h (scale bar, 50 μm, n = 3), respectively. (**I**) Flow cytometry analysis of cell cycle in differentiated HPCs cultured with or without 5 mM fructose in the presence or absence of 10 ng/mL recombinant human IL-6 for 72 h (n = 6), respectively. (**J**) Flow cytometry analysis of apoptosis in differentiated HPCs cultured with or without 5mM fructose in the presence or absence of 10 ng/mL recombinant human IL-6 for 72 h (n = 6), respectively. (**K**) Immunofluorescence analysis of EDU (green) in differentiated HPCs transfected with *TTP* siRNA or its control, and then incubated with or without 5 mM fructose for 72 h (scale bar, 50 μm, n = 3), respectively. (**L**) Flow cytometry analysis of cell cycle in differentiated HPCs transfected with *TTP* siRNA or control siRNA, and then incubated with or without 5 mM fructose for 72 h (n = 6), respectively. (**M**) Flow cytometry analysis of apoptosis in differentiated HPCs transfected with *TTP* siRNA or control siRNA, and then incubated with or without 5 mM fructose for 72 h (n = 6), respectively. All data are expressed as mean ± SEM. * *p* < 0.05, ** *p* < 0.01, *** *p* < 0.001 compared with normal animal control group or normal cell control.

### Anti-inflammation attenuates fructose-induced podocyte hypertrophy

Maslinic acid abolished fructose-induced high expression of miR-92a-3p and down-regulation of P57 (Fig. 5A-B) to improve podocyte overproliferation (Fig. 5C-D) and apoptosis (Fig. 5E). The down-regulation of podocyte nephrin and podocin induced by fructose was significantly ameliorated by maslinic acid (Fig. 5F). As a result, podocyte hypertrophy in fructose-exposed HPCs (Fig. 5G) was significantly improved by maslinic acid. These observations further demonstrated that podocyte hypertrophy induced by fructose could be alleviated by IL-6 reduction.

**Fig.5.**
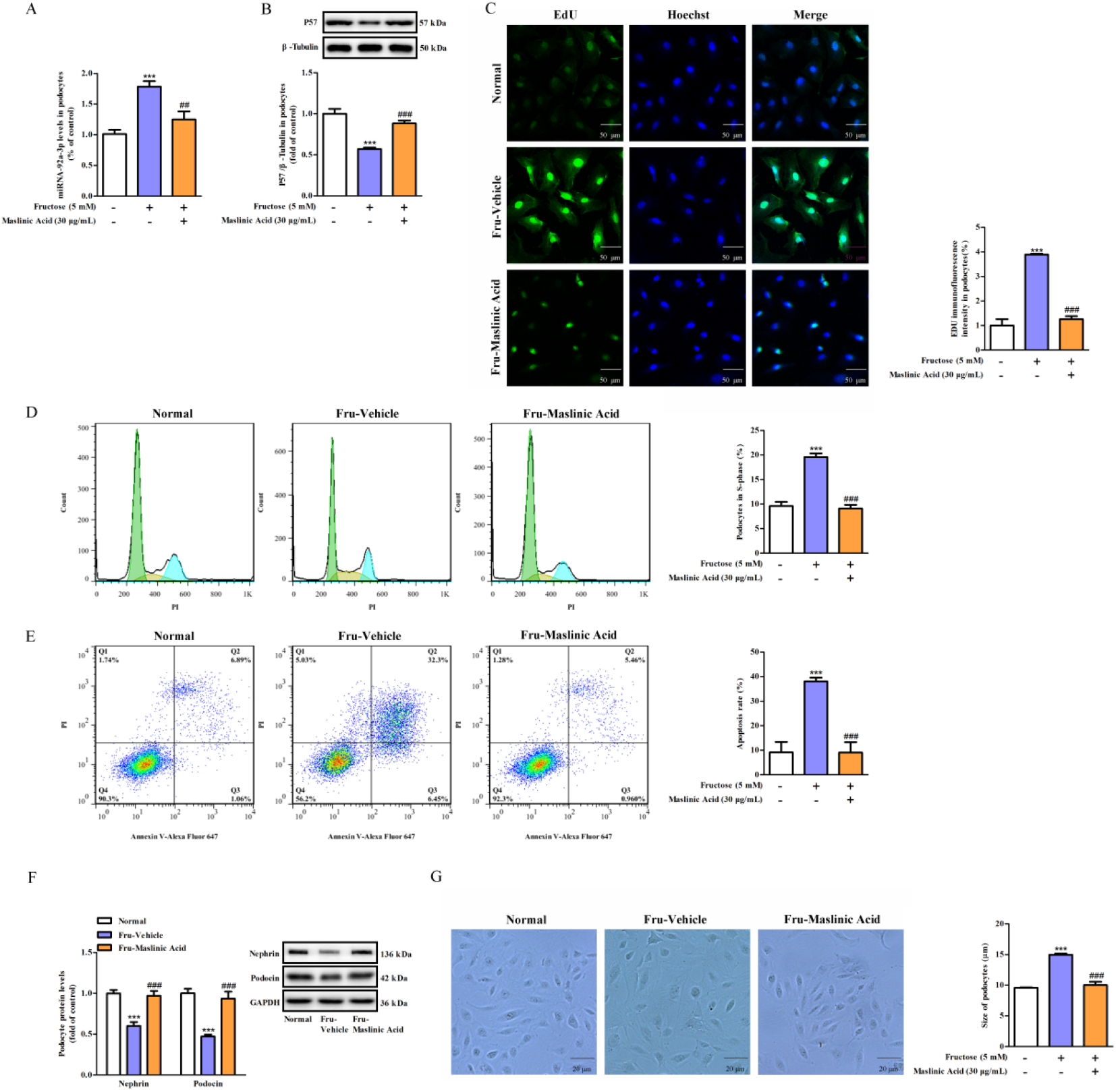
Maslinic acid abolishes fructose-induced podocyte proliferation and apoptosis in differentiated HPCs. (**A**) qRT-PCR analysis of miR-92a-3p levels in differentiated HPCs incubated with or without 5 mM fructose in the presence or absence of 30 μg/mL maslinic acid for 72 h (n = 6), respectively. (**B**) Western blot analysis of P57 protein levels in differentiated HPCs incubated with or without 5 mM fructose in the presence or absence of 30 μg/mL maslinic acid for 72 h. Relative protein level of P57 was normalized to β-Tubulin (n = 6), respectively. (**C**) Immunofluorescence analysis of EDU (green) in differentiated HPCs cultured with or without 5 mM fructose in the presence or absence of 30 μg/mL maslinic acid for 72 h (scale bar, 50 μm, n = 3), respectively. (**D**) Flow cytometry analysis of cell cycle in differentiated HPCs cultured with or without 5 mM fructose in the presence or absence of 30 μg/mL maslinic acid for 72 h (n = 6), respectively. (**E**) Flow cytometry analysis of apoptosis in differentiated HPCs cultured with or without 5 mM fructose in the presence or absence of 30 μg/mL maslinic acid for 72 h (n = 6), respectively. (**F**) Western blot analysis of nephrin and podocin protein levels in differentiated HPCs incubated with or without 5 mM fructose in the presence or absence of 30 μg/mL maslinic acid for 72 h. Relative protein levels of nephrin and podocin were normalized to GAPDH (n = 6), respectively. (**G**) Podocyte size in differentiated HPCs cultured with or without 5 mM fructose in the presence or absence of 30 μg/mL maslinic acid was measured separately for 72 h (scale bar, 20 μm, n = 3). All data are expressed as mean ± SEM. *** *p* < 0.001 compared with normal animal control group or normal cell control; ^##^*p* < 0.01, ^###^*p* < 0.001 compared with compared with fructose-vehicle animal group or fructose vehicle cell group.

### High fructose up-regulates KHK-A expression and restrains TTP expression to activate IL-6/STAT3 signaling in podocyte hypertrophy

Fructose increased KHK activity in differentiated HPCs with fructose-dependent ATP depletion (Fig. 6A). In addition, up-regulation of KHK-A protein levels (Fig. 6B-C) were detected in the glomeruli of fructose-fed rats and fructose-exposed differentiated HPCs. The correlation between high fructose-induced KHK-A dysregulation and podocyte injury were analyzed. QRT-PCR and Western blot analysis confirmed the high transfection efficiency of *KHK-A* siRNA in differentiated HPCs (Fig. 6D-E). *KHK-A* siRNA abolished fructose-induced down-regulation of TTP protein levels, overproduction of IL-6 and phosphorylation of STAT3 in differentiated HPCs (Fig. 6F-H). Immunoprecipitation results revealed an interaction between KHK-A and TTP (Fig. 6I). Furthermore, *KHK-A* siRNA reduced EdU-positive podocytes and the proportion of podocytes in S phase in fructose-exposed differentiated HPCs (Fig. 6J-K). Moreover, *KHK-A* siRNA improved apoptosis (Fig. 6L) and hypertrophy (Fig. 6M) induced by fructose exposure in differentiated HPCs. These observations showed that blockage of KHK-A may up-regulate TTP to inhibit the podocyte overproliferation, apoptosis and hypertrophy via activating of IL-6/STAT3 signaling.

**Fig.6.**
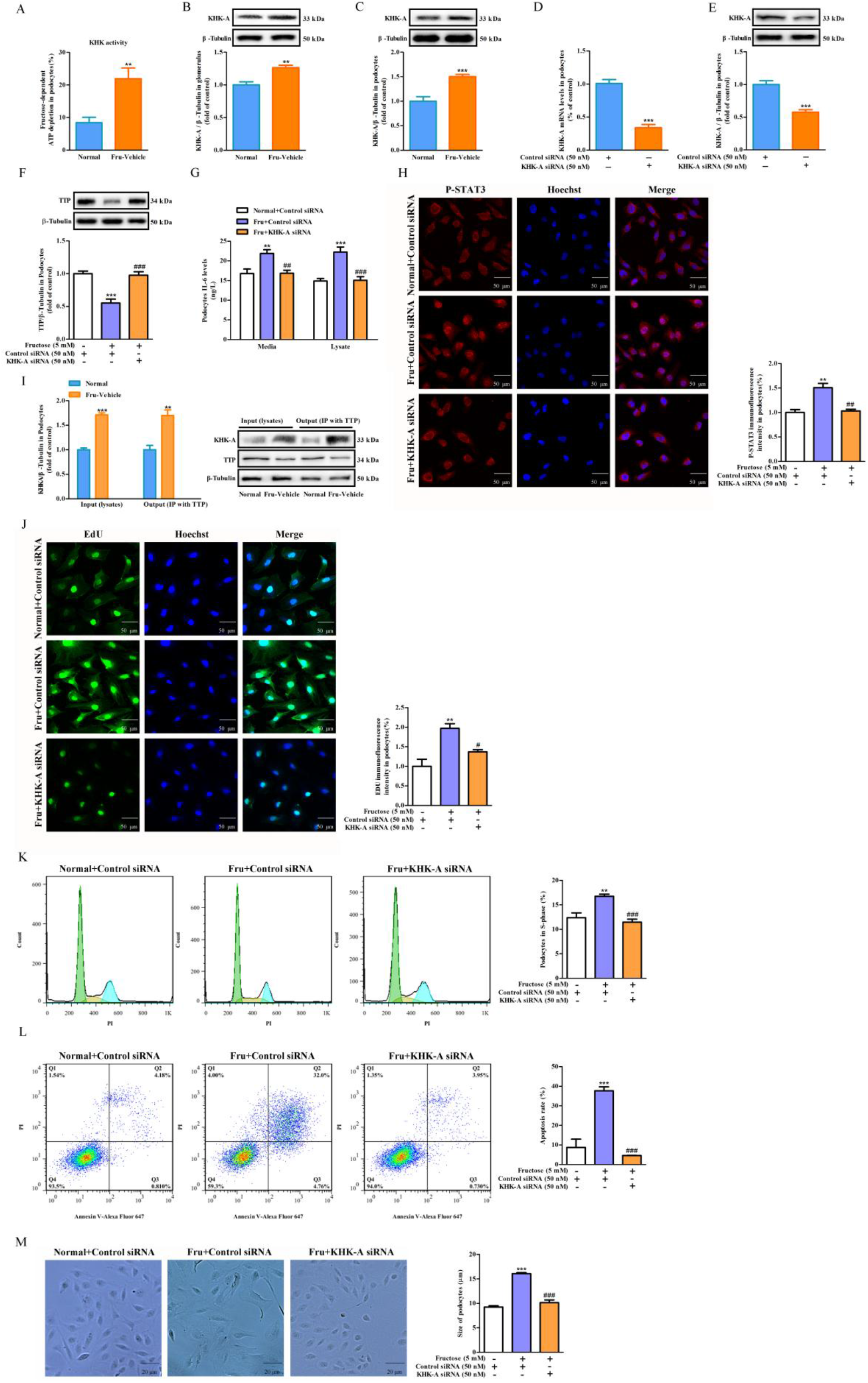
Transfection with KHK-A siRNA down-regulates IL-6 levels to alleviate fructose-induced podocyte proliferation, apoptosis and hypertrophy. (**A**) Enzyme activity test of KHK in differentiated HPCs (n = 8) of normal control, fructose vehicle groups, cultured with or without 5 mM fructose for 72 h, respectively. (**B-C**) Western blot analysis of KHK-A protein levels in the kidney glomeruli (16W) of normal control, fructose vehicle rats or differentiated HPCs cultured with or without 5 mM fructose for 72 h. Relative protein level of KHK-A was normalized to β-Tubulin (n = 6), respectively. (**D-E**) The transfection efficiency of KHK-A gene silencing in differentiated HPCs transfected with *KHK-A* siRNA (50 nM) or control siRNA was evaluated by qRT-PCR analysis (24 h, n = 6), and Western blot analysis (48 h, n = 6), respectively. (**F**) Western blot analysis of TTP (n = 6) protein levels in differentiated HPCs transfected with *KHK-A* siRNA or control siRNA, and then incubated with or without 5 mM fructose for 72 h, respectively. (**G**) ELISA analysis of IL-6 (n = 6) expression from media and lysate in differentiated HPCs transfected with *KHK-A* siRNA or control siRNA, and then incubated with or without 5 mM fructose for 72 h, respectively. (**H**) Immunofluorescence analysis of P-STAT3 (red) in differentiated HPCs transfected with *KHK-A* siRNA or control siRNA, and then incubated with or without 5 mM fructose for 72 h, (scale bar, 50 μm, n = 3), respectively. (**I**) Immunoprecipitation analysis of KHK-A and TTP protein interaction in differentiated HPCs incubated with or without 5 mM fructose for 72 h (n = 6). (**J**) Immunofluorescence analysis of EDU (green) in differentiated HPCs transfected with *KHK-A* siRNA or control siRNA, and then incubated with or without 5 mM fructose for 72 h (scale bar, 50 μm, n = 3), respectively. (**K**) Flow cytometry analysis of cell cycle in differentiated HPCs transfected with *KHK-A* siRNA or control siRNA, and then incubated with or without 5 mM fructose for 72 h (n = 6), respectively. (**L**) Flow cytometry analysis of apoptosis in differentiated HPCs transfected with *KHK-A* siRNA or control siRNA, and then incubated with or without 5 mM fructose for 72 h (n = 6), respectively. (**M**) Podocyte size in differentiated HPCs transfected with *KHK-A* siRNA or control siRNA, and then incubated with or without 5 mM fructose for 72 h was measured separately (scale bar, 20 μm, n = 3). All data are expressed as mean ± SEM. ** *p* < 0.01, *** *p* < 0.001 compared with normal animal control group or normal cell control; ^#^*p* < 0.05, ^##^*p* < 0.01, ^###^*p* < 0.001 compared with compared with fructose-vehicle animal group or fructose vehicle cell group.

## Discussion

High fructose induces kidney enlargement and cell overproliferation of renal cortex in rats (Sánchez-Lozada et al., 2007; Nakayama et al., 2010), however, the underlying mechanism remains unclear. In this study, we proved that the activation of IL-6/STAT3 signaling was the central event in high fructose-induced glomerular and podocyte hypertrophy. And it is firstly manifested that KHK-A-mediated TTP down-regulation negatively regulated IL-6/STAT3 signaling in podocyte hypertrophy under fructose exposure.

In this work, we observed IL-6/STAT3 signaling activation in hypertrophic glomeruli of fructose-fed rats and hypertrophic podocyte under fructose exposure. Animal-free recombinant human IL-6 increased STAT3 phosphorylation and podocyte hypertrophy in differentiated HPCs, which were consistent with other report in cultured mouse podocytes (Jo et al., 2016). Maslinic acid is a natural pentacyclic triterpene compound with anti-inflammatory effect (Lozano-Mena et al., 2014). Maslinic acid is able to inhibit cell proliferation and STAT3 phosphorylation by down-regulating IL-6 expression in human gastric cancer cells (Wang et al., 2017). In this study, this compound was found to suppress IL-6/STAT3 signaling activation, resulting in its attenuation of fructose-induced podocyte hypertrophy.

TTP has been reported to be a negative regulator of inflammatory cytokine IL-6 (Guo et al., 2020; Zhang et al., 2020). Its low expression was detected in glomeruli of fructose-fed rats and fructose-exposed HPCs, which were consistent with that in glomeruli of diabetic kidney disease patients (Guo et al., 2020). TTP overexpression obviously decreases IL-6 levels in human proximal tubular epithelial cells (Zhang et al., 2020). Its knockdown enhances podocyte injury marker claudin-1 expression and inflammatory response (Guo et al., 2020). Here, *TTP* siRNA interference did not further activate fructose-induced IL-6/STAT3 signaling and podocyte hypertrophy in HPCs, showing that the negative regulatory effect of TTP on IL-6/STAT3 signaling may be sensitive to fructose exposure in podocyte hypertrophy. Thus, the activation of IL-6/STAT3 signaling by TTP blockage may be an essential hallmark in fructose-induced podocyte hypertrophy.

Our previous study showed inflammatory response and apoptosis in glomerular podocytes of fructose-fed rats (Wang et al., 2015; Li et al., 2019). Podocyte proliferation was also detected in the glomeruli of fructose-fed rats and fructose-exposed differentiated HPCs. These observations led us to further explore podocyte proliferation and apoptosis in fructose-induced IL-6/STAT3 signaling activation and podocyte hypertrophy. P57, a cyclin-dependent kinase inhibitor, is reported to be constitutively expressed in mature podocytes (Nagata et al., 1998; Shankland and Wolf, 2000). Low expression of P57 was also observed in rat glomeruli and differentiated HPCs under high fructose, which were consistent with other report in the glomeruli of collapsing glomerulopathy patients with podocyte overproliferation (Barisoni et al., 2000). Podocyte proliferation is limited in primary cultured podocytes transfected with anti-miR-92a compared with those cells transfected with anti-miR control (Henique et al., 2017). We found that high fructose increased miR-92a-3p expression in rat glomeruli and differentiated HPCs. Based on the analysis from TargetScan in this study, P57 showed a 3’untranslated region binding sequence of miR-92a-3p, indicating that miR-92a-3p could directly targeted P57. These data indicated that overexpression of miR-92a-3p and down-regulation of P57 were closely related to podocyte proliferation and apoptosis in fructose-induced podocyte hypertrophy.

Furthermore, we showed that animal-free recombinant human IL-6 caused overexpression of miR-92a-3p and down-regulation of P57, as well as podocyte proliferation and apoptosis in differentiated HPCs. However, *TTP* siRNA interference did not further induce podocyte proliferation and apoptosis in fructose-exposed differentiated HPCs. Of note, maslinic acid significantly blocked the increased phosphorylation of STAT3, overexpression of miR-92a-3p and down-regulation of P57 in fructose-exposed differentiated HPCs. As a result, podocyte overproliferation, apoptosis and hypertrophy induced by fructose were significantly ameliorated. Therefore, we postulated that the activation of IL-6/STAT3 signaling mediated podocyte proliferation and apoptosis, contributing to fructose-induced podocyte hypertrophy. Inhibition of IL-6/STAT3 activation may alleviate podocyte proliferation, apoptosis and hypertrophy.

KHK-A catalyzes fructose to undergo the first catabolic biochemical reaction, and its dysregulation draws forth the multiple metabolic disturbances induced by high fructose consumption (Lanaspa et al., 2014). In the adipose tissue from KHK-A/C-KO mice, the overexpression of monocyte chemoattractant protein-1 and tumor necrosis factor-α induced by fructose are completely prevented, demonstrating that fructose-induced inflammation is dependent on KHK-A (Marek et al., 2015 Feb). Fructose-induced up-regulation of intracellular fructose-1-phosphate content is significantly inhibited by *KHK* siRNA in porcine renal proximal epithelial cells (Hayasaki et al., 2019). Knockdown of KHK abolishes fructose-induced monocyte chemoattractant protein-1 overproduction in human proximal tubular epithelial cells (Cirillo et al., 2009). We observed high KHK-A protein levels in the glomeruli of fructose-fed rats and fructose-exposed differentiated HPCs. As a result, ATP depletion induced by fructose was detected in differentiated HPCs. Immunoprecipitation results revealed that overexpression of KHK-A was closely correlated with low-expression of TTP in fructose-exposed differentiated HPCs. Furthermore, *KHK-A* siRNA significantly ameliorated fructose-induced TTP down-regulation in differentiated HPCs. More importantly, activation of IL-6/STAT3 signaling, podocyte overproliferation, apoptosis and hypertrophy induced by fructose were mitigated in differentiated HPCs transfected with *KHK-A* siRNA. These observations indicated that fructose-induced podocyte hypertrophy may probably result from KHK-A overexpression and TTP down-regulation. Consequently, the activation of IL-6/STAT3 signaling may trigger overproliferation and apoptosis, leading to fructose-induced podocyte hypertrophy.

In summary, this study demonstrated that high fructose increased KHK-A to trigger TTP down-regulation, subsequently activate IL-6/STAT3 signaling to increase miR-92a-3p and decrease its target protein P57 levels, in which, podocyte underwent overproliferation and apoptosis, finally causing hypertrophy. The inactivation of IL-6/STAT3 signaling by anti-inflammation may be a novel strategy for the treatment of podocyte hypertrophy in high fructose-associated kidney diseases.

## Materials and Methods

### Animals and treatment

All animal experiments were approved by the Institutional Animal Care and Use Committee of Nanjing University. Male Sprague-Dawley rats (5 weeks old, 200-220 g body weight) were purchased from the Experimental Animal Center of Zhejiang province (Hangzhou, China). The animals were housed with water and food *ad libitum* in a temperature and humidity controlled room with a 12:12 h light-dark cycle. 12 rats were divided into two experimental groups: normal group (n = 6) had standard diet and drinking water (available *ad libitum*), and fructose-fed group (n = 6) had standard diet and 10% fructose solution in drinking water (wt/vol, available *ad libitum*). After 8 weeks, high fructose-fed group still received 10% fructose in drinking water. Rats in normal group received saline solution for 8 weeks as the control. Body weight was recorded once a week throughout the experiment.

At week 15, each rat was placed in a metabolic cage to collect 24 h urine, respectively. Urine samples were centrifuged at 3,000 rpm for 10 min at 4 °C to remove particulate contaminants and stored at −80 °C. All efforts were made to minimize animal suffering and reduce the number of animals used.

### Reagents

For animal experiments, fructose was provided from Shandong Xiwang Sager Industry Co., Ltd. (Binzhou, China). For cell experiments, fructose was purchased from Sigma-Aldrich Inc. (St. Louis, MO). RPMI-1640 and opti-MEM were purchased from Basal Media Biotechnology Co., Ltd (Shanghai, China). Lipofectamine 2000, Trizol reagent, Hoechst 33342 (H3570), Alexa Fluor 488 goat anti-rabbit IgG (A11008) and Alexa Fluor 555 goat anti-rabbit IgG (A21428) were purchased from Invitrogen Biotechnology Co., Ltd (Shanghai, China). Fetal bovine serum (FBS) was purchased from Excell BioCorporation (Wellington, New Zealand). Recombinant IFN-γ, HRP-conjugated mouse anti-IgG (HAF007) and HRP-conjugated goat anti-IgG (HAF017) were purchased from R&D Systems (Minneapolis, USA). Urine protein test kit (C035-2-1), creatinine assay kit (C011-2-1), urea assay kit (C013-2-1) and uric acid test kit (C012-2-1) were purchased from Jiancheng Biotechnology Co., Ltd (Nanjing, China). HiScript^®^ II One Step qRT-PCR Probe Kit, ChamQ™ SYBR^®^ qPCR Master Mix (Without ROX), M-MLV (H-) Reverse Transcriptase, dNTPs and RNase inhibitor were got from Vazyme Biotechnology Co., Ltd (Nanjing, China). Cell lysis RIPA buffer, 4, 6-diamidino-2-phenylindole (DAPI) staining solution, phenylmethanesulfonyl fluoride (PMSF), BeyoClick™ EdU Cell Proliferation Kit with Alexa Fluor 488 (C0071S), Cell Cycle and Apoptosis Analysis Kit (C1052) and ATP Assay Kit (S0026B) were purchased from Beyotime Biotechnology (Nanjing, China). Pierce™ BCA protein assay kit was purchased from Thermo Scientific (Schwerte, Germany). Rat IL-6 ELISA kit (YFXER00062) and human IL-6 ELISA kit (YFXEH00258) were purchased from YIFEIXUE Biotechnology Co., Ltd (Nanjing, China). Annexin V-PE/7-AAD apoptosis kit was purchased from FCMACS Biosciences Co., Ltd (Nanjing, China). Rabbit anti-nephrin (ab58968) and Rabbit anti-NPHS2 (ab50339) were purchased from Abcam (Cambridge, MA, USA). Rabbit ZFP36 antibody (12737-1-AP), anti-β-tubulin (10068-1-AP), Animal-free recombinant human IL-6 (HZ-1019), Immunoprecipitation Kit (PK10007), HRP-conjugated Goat Anti-Mouse IgG (SA00001-1) and HRP-conjugated Goat Anti-Rabbit IgG (SA00001-2) were purchased from Proteintech Group, Inc. (Chicago, USA). Rabbit anti-P-STAT3 (Tyr705) (D3A7) and Rabbit anti-STAT3α (D1A5) were purchased from Cell Signaling Technology (Cambridge, USA). Rabbit KHK Isoform A Antibody (21708) was purchased from Signalway Antibody LLC (Maryland, USA). Mouse anti-P57 (sc-56341) and Rabbit anti-GAPDH (sc-25778) were purchased from Santa Cruz Biotechnology Co., Ltd (Santa Cruz, CA, USA). Mouse anti-Synaptopodin (MAB4919) was purchased from Abnova Corporation (Taipei, Taiwan). Maslinic acid was purchased from APExBIO Technology LLC (Houston, USA).

### Blood and tissue processing

After the animal experiment, rats were anesthetized intraperitoneally using 50 mg/kg sodium pentobarbital after a 12 h-fast. Blood samples were collected from rat carotid artery and then centrifuged at 3,000 rpm for 10 min to get the serum. Following blood collection, rat kidney cortex was cut into equal pieces for glomeruli isolation and histology analyses, as well as total RNA isolation and protein isolation, respectively. Glomeruli were isolated by a graded sieving technique using sieves with pore sizes of 250, 150, and 75 μm as described previously (Wang et al., 2015). Briefly, rat kidney cortex tissues were cut into small pieces and pressed through 250- and 150-μm stainless steel mesh and then thoroughly washed with PBS on 75-μm stainless meshes. The rat kidney cortex tissues were fixed with 4% paraformaldehyde. The serum and glomerulus samples were then stored at −80 °C for further analyses.

### Biochemical analysis

Serum levels of uric acid, creatinine, urea nitrogen and proteinuria were detected by common commercially available biochemical kits. IL-6 level in the serum and renal cortex of rat, in the media and lysate of differentiated HPCs was detected by the commercially available ELISA kits. All experimental operations were standardized and in accordance with the manufacturer’s instructions.

### Cell culture and treatment

Heat-sensitive human podocyte cell line was obtained from Dr. Zhi-Hong Liu (Research Institute of Nephrology, Nanjing General Hospital of Nanjing Military Command, Nanjing, China), which was provided by M. Saleem (University of Bristol, Bristol, United Kingdom) (Li et al., 2019). The cells were cultivated at the permissive temperature (33 °C) in RPMI-1640 medium supplemented with 10% fetal bovine serum and recombinant IFN-γ.

To induce differentiation, these HPCs were seeded in 6-well plastic culture plates at a density of 5×10^4^ cells/mL under non-per-missive condition at 37 °C for 7 days in the absence of interferon-γ. Cell culture was replaced with fresh medium as appropriate. Differentiated HPCs were then cultured in RPMI-1640 medium (containing 10% fetal bovine serum) supplemented with or without 5 mM fructose to detect miR-92a-3p expression and IL-6 mRNA level by qRT-PCR analysis, the protein levels of KHK-A, TTP, P-STAT3, P57, podocin and nephrin by western blot analysis, P-STAT3 and EDU by immunofluorescence analysis, podocyte apoptosis and cell cycle by Flow cytometry analysis, podocyte size by microscopy analysis, respectively. The concentration of fructose was selected based on our previous studies and other reports (Cirillo et al., 2009; Wang et al., 2015).

Differentiated HPCs were incubated in RPMI-1640 medium (containing 10% fetal bovine serum) with or without 5mM fructose in the presence or absence of 30 μg/mL maslinic acid or 10 ng/mL animal-free recombinant human IL-6 in 6-well plates for 72 h to assay P-STAT3, P57, podocin, nephrin levels by Western blot analysis, miR-92a-3p expression by qRT-PCR, EDU and P-STAT3 by immunofluorescence, podocyte apoptosis and cell cycle by Flow cytometry, podocyte size by microscopy.

*TTP* siRNA (Biotend, Shanghai, China), *KHK-A* siRNA (GenePharma, Shanghai, China) as well as the respective negative control siRNA were synthesized respectively. The nucleotide sequences were listed in Table 1. *TTP* siRNA, *KHK-A* siRNA as well as the respective negative control were transiently transfected into the cultured differentiated HPCs with Lipofectamine 2000, according to the manufacturer’s instructions, respectively. The efficiency of transfection was evaluated by measuring TTP and KHK-A mRNA levels at 24 h by quantitative real-time PCR analysis, as well as TTP and KHK-A protein levels at 48 h by Western blot analysis, respectively. After 6 h, *TTP* siRNA or *KHK-A* siRNA transfected differentiated HPCs were incubated in RPMI-1640 medium (containing 10% fetal bovine serum) with or without 5 mM fructose for 72 h to test IL-6 by ELISA, EDU and P-STAT3 by immunofluorescence, podocyte cell cycle and apoptosis by Flow cytometry, podocyte size by microscopy, respectively.

**Table. 1.**
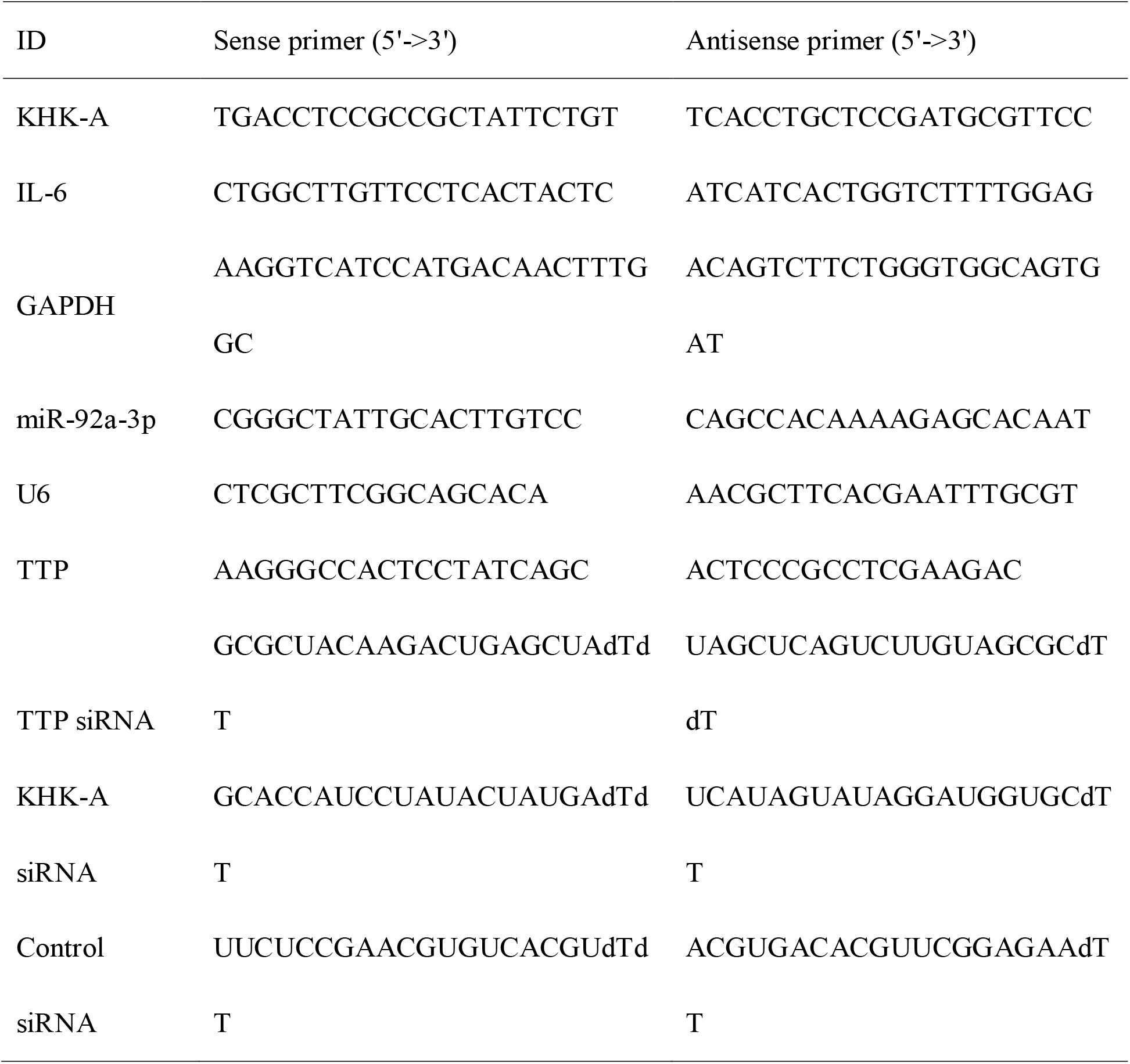
Primer sequences used for miRNAs and qRT-PCR in this study

**Table. 2.** Body weight and renal function damage index in fructose fed rats.

| Group | Body weight (g) | Serum uric acid<br>($\mu\text{mol/L}$) | Serum creatinine<br>($\mu\text{mol/L}$) | Serum urea<br>nitrogen (mmol/L) |
| --- | --- | --- | --- | --- |
| Normal | 496.33 ± 100.51 | 85 ± 3.24 | 93.42 ± 7.3 | 1.99 ± 0.27 |
| Fru-Vehicle | 838.43 ± 85.82<sup>***</sup> | 124.11 ± 2.91<sup>***</sup> | 159.31 ± 12.01<sup>***</sup> | 3.5 ± 0.13<sup>***</sup> |

Quantitative data are presented as mean ± SEM for n = 6 rats per group. <sup>\*\*\*</sup> $p < 0.001$ versus normal animal control group, there were no significant differences in all indicators in fructose-fed group compared with normal control group.

In all experiments, total cellular proteins and total RNAs were extracted, respectively. These samples were stored at −80 °C before Western blot or qRT-PCR assay. The protein concentration was determined by BCA protein assay kit.

### Immunofluorescence assay

The tested HPCs were washed with PBS for three times and fixed with 4% paraformaldehyde for 30 min. These cells were blocked with immunostainings blocking buffer containing 0.5% Triton X-100 for 40 min and washed by PBS. Then, the HPCs were labeled with primary antibody rabbit against P-STAT3 (1:100) at 4 °C overnight. P-STAT3 expression was detected with Alexa Fluor 555 goat anti-rabbit (dilution, 1:500) secondary antibody. The nuclei were counterstained with Hoechst 33342 and coverslip with aqueous mounting medium. Finally, all samples were examined with a confocal laser scanning microscope (Lei TCS SP8-MaiTai MP; Leica, Wetzlar, Germany).

Differentiated HPCs, cultured in RPMI-1640 medium (containing 10% fetal bovine serum) supplemented with or without 5 mM fructose for 72 h in the presence or absence of maslinic acid, animal-free recombinant human IL-6, transfected with *TTP* siRNA or *KHK-A* siRNA and corresponding controls. These cells were incubated with 10 μM EdU solution for 4 h according to instructions of EdU assay kit. Briefly, cells were fixed, permeabilized and cultured in click additive solution for 30 min. Then, the cell nucleus was stained with Hoechst 33342 solution. The results were observed and captured using confocal laser scanning microscope (Lei TCS SP8-MaiTai MP; Leica, Wetzlar, Germany).

Immunohistofluorescence staining was performed to detect the expression of nephrin and podocin proteins in rat glomeruli. Rat kidney cortex tissues were fixed with 4% paraformaldehyde, embedded in paraffin, and sectioned transversely. Thin frozen section (10 μm) of cortex tissues was blocked for 1 h with immunolstaining blocking buffer. The primary antibodies against nephrin (1:100), podocin (1:100) and KI67 (1:100) were used for incubation overnight at 4 °C. After washing with 0.01 M phosphate buffered solution (PBS) for three times, the sections were incubated with Alexa Fluor 555 goat anti-rabbit (1:500) or Alexa Fluor 488 goat anti-rabbit (1:500) secondary antibodies for 60 min at room temperature, and then stained with DAPI for 15 min. Finally, the sections were examined by confocal laser scanning microscope (Lei TCS SP8-MaiTai MP; Leica, Wetzlar, Germany).

### Tunnel assay

Rat renal cortex samples were fixed with 4% paraformaldehyde, embedded in paraffin, and sectioned transversely. The embedded sections were stained by the TUNEL technique using an in situ apoptosis detection kit according to the protocol provided by the manufacturer. Then, the nuclei were counterstained with DAPI. The apoptotic cells with green fluorescence in rat glomeruli were detected with a confocal laser scanning microscope (Lei TCS SP8-MaiTai MP; Leica, Wetzlar, Germany).

### PAS analysis

Rat renal cortex samples were fixed with 4% paraformaldehyde, embedded in paraffin, and sectioned transversely. The sections (10 μm) were subjected to PAS staining and were detected with a confocal laser scanning microscope (Lei TCS SP8-MaiTai MP; Leica, Wetzlar, Germany).

### Podocyte apoptosis and cell cycle analysis

Differentiated HPCs, cultured in RPMI-1640 medium (containing 10% fetal bovine serum) supplemented with or without 5 mM fructose in the presence or absence of maslinic acid, animal-free recombinant human IL-6, transfected with *TTP* siRNA or *KHK-A* siRNA and corresponding controls were harvested after 72 h incubation. Cells were washed with cold 0.01 M PBS for three times after the digestion with trypsin (without EDTA). The cell concentration was adjusted to 1× 10^6^ cells/mL. Cells were fixed and stained with following the protocol of Annexin V-PE/7-AAD apoptosis kit or the cell cycle analysis kit. Data was acquired by FACScan flow cytometer (BD Biosciences, Franklin Lakes, USA) and analyzed with Flowjo software.

### Podocytes morphological analysis

Differentiated HPCs were cultured in RPMI-1640 medium (containing 10% fetal bovine serum) supplemented with or without 5 mM fructose in the presence or absence of maslinic acid, animal-free recombinant human IL-6, transfected with *TTP* siRNA or *KHK-A* siRNA and corresponding controls for 72 h. The podocytes size was observed under inverted microscope (OLYMPUS, Tokyo, Japan) and quantified via ImageJ (version 1.42q, National Institutes of Health).

### KHK activity detection

Differentiated HPCs cultured in RPMI-1640 medium (containing 10% fetal bovine serum) supplemented with or without 5 mM fructose for 72 h were harvested in PBS. These cells were broken by ultrasound and then centrifuged at 15,000 rpm for 10 min at 4 °C to obtain cell lysate. Cell lysate was quantified to 5 mg/mL by BCA protein assay kit. For KHK enzyme activity analysis, the enzyme activity detection reaction included 50 μL of 5mg/mL cell lysate, 50 μL of PBS, 50 μL of 5 mM ATP, 50 μL of 5 mM fructose or ddH_2_O as control. The reaction mixture was incubated for 2 h at 37 °C. After deproteinization with perchloric acid and potassium hydroxide, the reaction mixture was prepared to detect ATP content by ATP assay kit according to the protocol provided by the manufacturer (Lanaspa et al., 2014). Date was evaluated by Luminometer protocol using CLARIOstar (BMG LABTECH, Aufenburg, Germany). KHK activity was calculated as the ratio between ATP level versus baseline for each sample and values were compared between normal and fructose vehicle groups.

### qRT-PCR analysis

Total RNA was isolated from rat kidney glomeruli and differentiated HPCs using trizol reagent according to the manufacturer’s recommendations, respectively. The primers used were listed in Table 1. These primers were synthesized by Generay Biotechnology Co., Ltd. (Shanghai, China). The reverse transcription reaction of mRNA used the HiScript^®^ II One Step qRT-PCR Probe Kit. The reverse transcription reaction of miRNAs used the M-MLV (H-) Reverse Transcriptase. The qRT-PCR analysis was performed in triplicate with ChamQ™ SYBR^®^ qPCR Master Mix (Without ROX) using the CFX96 Real-Time PCR Detection System (Bio-Rad), respectively. Gene expression was normalized to the level of GAPDH for mRNA and U6 for miRNA. Relative expressions of target genes were determined by the Ct(2^-ΔΔCt^) method.

### Western blot analysis

Proteins extracted from rat glomeruli and differentiated HPCs were separated by SDS-PAGE, and the proteins were electrophoretically transferred to polyvinylidene fluoridemembranes. Nonspecific protein-binding sites were blocked with TBS containing 0.1% Tween-20 and 5% non-fat milk for 1 h at room temperature, and the blots were incubated with specific primary antibodies including anti-nephrin (dilution 1:1000), anti-podocin (dilution 1:1000), anti-KHK-A (dilution 1:1000), anti-TTP (dilution 1:1000), anti-P-STAT3 (dilution 1:1000), anti-STAT3 (dilution 1:1000), anti-P57 (dilution 1:500), anti-β-Tubulin (dilution 1:1000) and anti-GAPDH (dilution 1:4000). Blots were incubated overnight at 4 °C, followed by HRP-conjugated anti-rabbit IgG antibody (dilution 1:20,000) or HRP-conjugated anti-mouse IgG antibody (dilution 1:20,000). Immunoreactive bands were visualized via enhanced chemiluminescence and quantified via densitometry using ImageJ (version 1.42q, National Institutes of Health).

### Immunoprecipitation analysis

Differentiated HPCs cultured in RPMI-1640 medium (containing 10% fetal bovine serum) supplemented with or without 5 mM fructose for 72 h were harvested in PBS. These cells were broken by ultrasound and then centrifuged at 15,000 rpm for 10 min at 4 °C to obtain cell lysate. Cell lysate was quantified to 5 mg/mL by BCA protein assay kit. The cell lysate was followed by Immunoprecipitation Kit according to the protocol provided by the manufacturer. Then, the samples obtained in the above step were analyzed by Western blot analysis. Immunoreactive bands were visualized via enhanced chemiluminescence and quantified via densitometry using ImageJ (version 1.42q, National Institutes of Health).

### Statistical analysis

All data were expressed as the mean ± SEM. For experiments with more than two subgroups, statistical comparisons were performed by a one-way analysis of variance, followed by Dunnett test. For experiments with two subgroups, statistical comparisons were performed by t tests by unpaired t test. In all statistical comparisons, *p* < 0.05 was considered to be significant. Figures were obtained by Graphpad Prizm Program 6.0 software (Graphpad, San Diego, CA).

## Acknowledgment

We thank Dr. Zhi-Hong Liu (Research Institute of Nephrology, Nanjing General Hospital of Nanjing Military Command, Nanjing, China) for providing Heat-sensitive human podocyte cell line. This work was supported by Grant from National Natural Science Foundation of China (No. 81730105) to Ling-Dong Kong.

## Conflict of interest

The authors declare no competing financial interests.

